# SAXS analysis of intrinsic tenase complex bound to lipid nanodisc highlights intermolecular contacts between factors VIIIa/IXa

**DOI:** 10.1101/2021.07.09.451823

**Authors:** Kenneth C. Childers, Shaun C. Peters, P. Lollar, H. Trent Spencer, Christopher B. Doering, P. Clint Spiegel

**Affiliations:** Department of Chemistry, Western Washington University, Bellingham, WA, USA; Department of Pediatrics, Aflac Cancer and Blood Disorders Center, Children’s Healthcare of Atlanta, Emory University, Atlanta, GA, USA

**Keywords:** factor VIII, factor IX, tenase, nanodiscs, small-angle x-ray scattering (SAXS)

## Abstract

The intrinsic tenase (Xase) complex, formed by factors (f)VIIIa and fIXa, forms on activated platelet surfaces and catalyzes the activation of factor X to Xa, stimulating thrombin production in the blood coagulation cascade. The structural organization of the membrane-bound Xase complex remains largely unknown, hindering our understanding of the structural underpinnings that guide Xase complex assembly. Here, we aimed to characterize the Xase complex bound to a lipid nanodisc with biolayer interferometry (BLI) and small angle X-ray scattering (SAXS). Using immobilized lipid nanodiscs, we measured binding rates and nanomolar affinities for fVIIIa, fIXa, and the Xase complex. An *ab initio* molecular envelope of the nanodisc-bound Xase complex allowed us to computationally model fVIIIa and fIXa docked onto a flexible lipid membrane and identify protein-protein interactions. Our results highlight multiple points of contact between fVIIIa and fIXa, including a novel interaction with fIXa at the fVIIIa A1-A3 domain interface. Lastly, we identified hemophilia A/B-related mutations with varying severities at the fVIIIa/fIXa interface that may regulate Xase complex assembly. Together, our results support the use of SAXS as an emergent tool to investigate the membrane-bound Xase complex and illustrate how mutations at the fVIIIa/fIXa dimer interface may disrupt or stabilize the activated enzyme complex.

## Introduction

Factor VIII (fVIII) is a procoagulant glycoprotein that is secreted into the bloodstream as a heterodimer with the domain architecture A1-A2-B/*a3*-A3-C1-C2 in a tight complex with von Willebrand factor (vWf) to inhibit premature fVIII clearance.^1,2^ Proteolytic cleavage by thrombin releases vWf and generates the activated fVIII (fVIIIa) A1/A2/A3-C1-C2 heterotrimer^3,4^ which binds to activated platelet surfaces through several hydrophobic loops in the C1 and C2 domains.^5–7^ Binding to activated factor IX (fIXa), a trypsin-like serine protease, forms the intrinsic tenase (Xase) complex, which catalyzes the activation of factor X (fX), enhancing thrombin turnover and clot formation.^8^ FIXa is a heterodimer of its light chain, composed of an N-terminal γ-carboxyglutamic acid (Gla) domain^9,10^ and two epidermal growth factor-like (EGF-1 and EGF-2) domains, and heavy chain, which carries the catalytic domain.^11^ Inactivation of the Xase complex occurs by dissociation of the fVIIIa A2 domain, proteolytic degradation by activated protein C, and inhibition of fIXa by antithrombin III.^12,13^

In the absence of fVIIIa and phospholipids, fIXa retains minimal activity due to the autoinhibitory 99-loop, which sterically blocks occupation of the active site and is stabilized by multiple intramolecular contacts.^8,14–16^ Binding to fVIIIa and fX on lipid membranes is hypothesized to rearrange the 99-loop, increasing *kcat* by ∼10^3^-fold and reducing *K*_*m*_ by ∼10^3^-fold.^17,18^ Individuals with hemophilia A or B, caused by a mutation to fVIII or fIX, respectively, have reduced capacity to form the Xase complex and significantly lowered clotting efficiency.^2,19^ Conversely, several gain-of-function mutations to fIX are proposed to stabilize the Xase complex and have been identified in patients with deep vein thrombosis,^20–22^ making the Xase complex a suitable target for treating thrombotic disorders.^23–26^

Structural studies on the Xase complex have proposed multiple binding sites between the fVIIIa A2, A3, and C2 domains, and fIXa catalytic and Gla domains.^27–33^ A putative allosteric network in the fIXa catalytic domain between the 99-loop and a cluster of solvent-exposed helices, collectively described as exosite II, has been proposed to modulate the conformation of the 99-loop^34,35^ and is a potential binding site for fVIIIa.^29,36^ The crystal structure of the prothrombinase complex, formed by factors Va/Xa, which are homologous to fVIIIa/fIXa, respectively, suggests the catalytic domain docks onto the A2 and A3 domains.^37^ Computational models of the Xase complex have provided broad structural analyses of how fVIIIa docks onto fIXa and promotes binding factor X.^38–40^ However, a lack of biochemical information on the complete lipid-bound Xase complex in solution hinders a mechanistic understanding of how the enzyme complex is formed.

In this study, we characterized the binding kinetics and solution structure of the Xase complex bound to a lipid nanodisc (Xase:ND). Using biolayer interferometry (BLI), we calculated nanomolar affinity between fVIIIa/fIXa and immobilized nanodiscs as well as association and dissociation rate constants. A molecular envelope was calculated through small-angle X-ray scattering (SAXS), allowing us to perform a combination of computational studies to determine a working model of the Xase:ND complex in solution. In addition to supporting previously proposed intermolecular contacts, our results identified a novel interaction between the fVIIIa A1/A3 domain interface and fIXa EGF-1 domain. These findings allowed us to speculate on hemophilia A and B-related mutations that may disrupt formation of the Xase complex.

## Materials and Methods

### Proteins

A bioengineered human/porcine chimera of B domain-deleted fVIII, termed ET3i,^41,42^ was activated using a Thrombin Cleavage Capture Kit (Millipore) according to the manufacturer’s specifications. EGRck-active site blocked human fIXa was purchased from Haematologic Technologies. The plasmid encoding for membrane scaffold protein MSP1D1 was purchased from AddGene.^43^

### Preparation of lipid nanodiscs

Lipid nanodiscs were assembled as previously described.^44–46^ Briefly, MSP1D1 was expressed in BL21(DE3) bacterial cells and purified using Ni^2+^-affinity resin. The His_6_-tagged protein was dialyzed into TBS (20 mM tris pH 8, 100 mM NaCl, 0.5 mM EDTA), concentrated to 10 mg/mL (400 µM), snap frozen in liquid N_2_, and stored at −80 °C. Lipids were prepared as an 80:20 molar ratio of 1,2-dioleoyl-sn-glycero-3-phosphocholine (DOPC) and 1,2-dioleoyl-sn-glycero-3-phospho-L-serine (DOPS) (Avanti Polar Lipids), respectively, at a working concentration of 6 mM in TBS supplemented with 100 mM cholate. MSP1D1 and lipids were mixed at a 1:47 molar ratio, respectively, with 20 mM cholate and incubated for 1 hour at room temperature. To initiate the assembly of lipid nanodiscs, 0.5 mL of activated BioBeads (BioRad) were added to the sample and incubated for 2.5 hours. BioBeads were removed using a 0.22 µm spin filter and nanodiscs were purified by size-exclusion chromatography (Superdex 200 Increase 10/300, GE Healthcare) with TBS as the running buffer. A single peak was obtained at 13 mL retention volume. Fractions were pooled, concentrated to 10 µM, and stored at 4 °C.^43,47^

### BLI measurements

Binding kinetics and affinities were measured on a BLItz (ForteBio) biosensor at room temperature. Anti-penta-HIS BLI tips (ForteBio) were activated in HBS (20 mM HEPES pH 7.4, 100 mM NaCl, 5 mM CaCl_2_) for 10 minutes and loaded with 4 µL of 400 nM His_6_-tagged nanodiscs for 60 seconds to establish a baseline. After washing in buffer, association and dissociation steps were observed over 90 seconds for fVIIIa, fIXa, and Xase (1:1, fVIIIa:fIXa) across a serial dilution (12.5-1600 nM). Binding curves were generated by GraphPad Prism 5.0 software (GraphPad Software, San Diego, CA) using a global fit analysis and 1:1 binding mode from an average of three or more independent BLI experiments to determine association (*k*_*on*_) and dissociation (*k*_*off*_) rates. The dissociation constant (*K*_*D*_) was defined as the ratio *k*_*off*_*/k*_*on*_.

### SAXS sample preparation and data collection

To form the Xase:ND complex, activated ET3i (5.3 µM) and fIXa (37.4 µM) were mixed with lipid nanodiscs (10 µM) at a 2:2:1 (fVIIIa:fIXa:ND) molar ratio. Samples of the empty nanodisc and nanodisc-bound Xase complex were buffer exchanged into HBS, supplemented with 10 mM KNO_3_ and 1% (w/v) sucrose to reduce radiation damage,^48^ using 3 kDa MWCO spin filters. BLI binding measurements confirmed that KNO_3_ and sucrose did not disrupt the Xase complex from binding to lipid nanodiscs (Figure S1). Both samples were concentrated to approximately 1 mg/mL (assuming an A_280_ of 1.0 is equivalent to a protein concentration of 1 mg/mL) and stored at 4 °C.

High-throughput SAXS data were collected on the Advanced Light Source SIBYLS beamline 12.3.1 at Lawrence Berkeley National Laboratory (Berkeley, CA). Scattering intensities for each sample were collected from concentrations of 0.33, 0.66, and 1.0 mg/mL every 0.3 seconds for a total of 33 images. To ensure binding between the lipid nanodisc and Xase complex during SAXS data collection, the molar concentration of Xase was maintained above the calculated *K*_*D*_ value. Buffer-subtracted intensities were averaged using FrameSlice (SIBYLS).

### Generation of ab initio envelopes and models

Scattering intensities were merged and processed with PRIMUS^49^ and GNOM^50^ to calculate Kratky, Guinier, and P(r) distribution plots for the empty nanodisc and Xase:ND complex. Six independent *ab initio* molecular envelopes were generated and averaged for each sample by DAMMIN^51^ and DAMAVER,^52^ respectively. A final molecular envelope was determined by DAMFILT^52^ with half the input volume as the molecular cutoff.

Solution structures of MSP1D1 (PDB ID: 6CLZ)^53^ and lipids (PDB ID: 2N5E)^54^ were used as templates for modeling the empty nanodisc with SREFLEX.^55^ To generate a model of the nanodisc-bound Xase complex, we performed a combination of computational docking, rigid body fitting, and flexible refinement. A complete structure of ET3i was generated by SWISS-MODEL^56^ to account for flexible loops absent in the crystal structure (PDB ID: 6MF0).^57^ Residues 1649-1689, comprising the acidic *a3* domain, were removed to produce a model of activated ET3i. RosettaDock^58–60^ allowed for initial docking experiments between activated ET3i and fIXa (PDB ID: 1PFX). Iterative rounds of SREFLEX^55^ accounted for flexibility during refinement for both models and the χ^2^ value was used as a metric to track refinement as calculated by FOXS.^61,62^

## Results

### Lipid binding characteristics of fVIIIa, fIXa, and Xase

Binding kinetics and affinities between fVIIIa, fIXa, and the Xase complex to immobilized lipid nanodiscs was measured with BLI. By retaining the His_6_-tag on the MSP1D1 scaffold protein, we loaded anti-His_6_ BLI tips with nanodiscs and measured association and dissociation rates with individual coagulation factors and the Xase complex (Figure 1, Table 1). FVIIIa displayed the highest affinity for lipid membranes with an apparent equilibrium dissociation constant (*K*_*D*_) of 32.3 ± 6.0 nM. Remarkably, fIXa had the lowest affinity, measuring a *K*_*D*_ of 199 ± 1.7 nM, largely due to a rapid dissociation rate of 0.0723 ± 0.003 s^-1^. We also confirmed that binding fIXa to lipid nanodiscs was Ca^2+^-dependent (Figure S2), as previously reported.^17^ Lastly, the Xase complex demonstrated high affinity for the immobilized nanodiscs, with a *K*_*D*_ of 71.3 ± 4.4 nM. To our knowledge, these data represent the first reported binding affinities and rate constants between activated tenase complex proteins and lipid nanodiscs.

**Table 1.** Apparent binding kinetics and affinities of fVIIIa, fIXa, and Xase with lipid nanodiscs. All data represent the average of three or more independent experiments with a 95% confidence interval. *k*_*on*_^-^ rate of association, *k*_*off*_^-^ rate of dissociation, *K*_*D*_^-^ dissociation constant (*k*_*off*_/*k*_*on*_).

| | $k_{on}$ ($\times 10^4 \text{ M}^{-1} \text{ s}^{-1}$) | $k_{off}$ ($\times 10^{-3} \text{ s}^{-1}$) | $K_D$ (nM) |
| --- | --- | --- | --- |
| fVIIIa | $8.62 \pm 0.11$ | $2.79 \pm 0.55$ | $32.3 \pm 6.0$ |
| fIXa | $36.3 \pm 1.2$ | $72.3 \pm 3.0$ | $199 \pm 1.7$ |
| Xase | $14.9 \pm 3.5$ | $10.7 \pm 0.90$ | $71.3 \pm 4.4$ |

**Figure 1.**
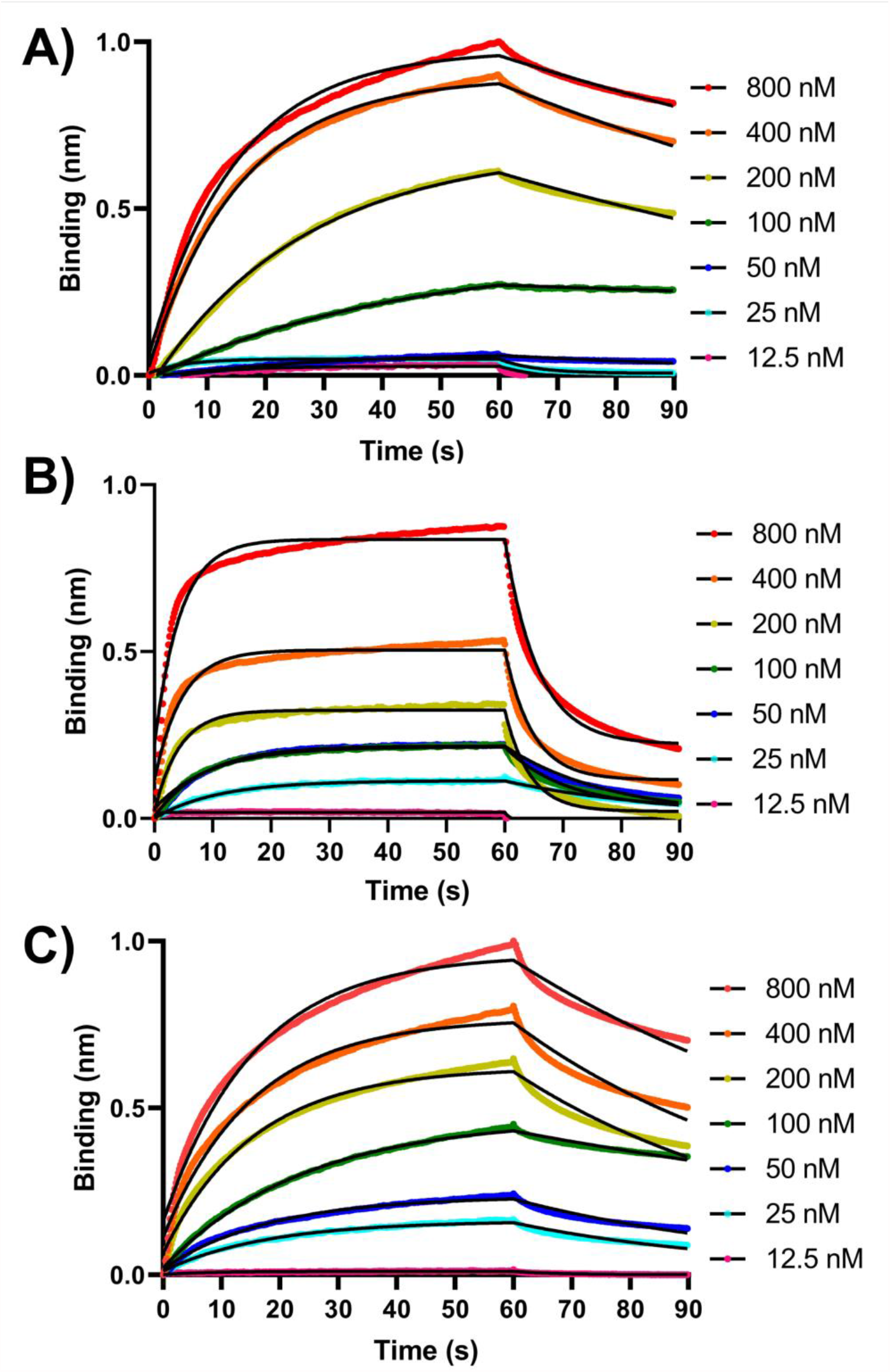
Bio-layer interferometry binding measurements. Apparent binding rates and affinities for **(A)** fVIIIa, **(B)** fIXa, and **(C)** Xase (1:1, fVIIIa:fIXa) were determined by biolayer interferometry (BLI) using immobilized His_6_-tagged lipid nanodiscs. Step-corrected association and dissociation steps were measured for 60 seconds and 30 seconds, respectively, over a serial dilution of each sample (spheres). The association and dissociation binding kinetics were calculated from a global fit using a 1:1 binding analysis (black line) (GraphPad Prism 5.0) and used to calculate *K*_*D*_ values (*k*_*off*_/*k*_*on*_). Results are summarized in Table 1.

### Solution structure of nanodisc

We next investigated the structural characteristics of the empty nanodisc in solution using SAXS. Based on our analysis, the unloaded nanodisc adopted a compact, globular structure in solution. (Figure 2A). Scattering intensities from the Guinier region of the scatter plot were used to calculate a radius of gyration (*Rg*) and radius of cross-section (*Rc*) of 54.9 Å and 33.6 Å, respectively (Figure 2B, 2C, Table 2). A final model of the empty nanodisc was generated through iterative rounds of refinement with SREFLEX and fit into the *ab initio* molecular envelope (Figure 2D). These findings indicate that the nanodisc adopts a bent, elliptical shape in solution, in good agreement with previously reported SAXS and SANS studies.^43,63–67^ Alignment of a theoretical SAXS curve for the modeled empty nanodisc with our experimental data yielded a χ^2^ value of 0.26, indicating strong agreement between the two sets of data.

**Table 2.**
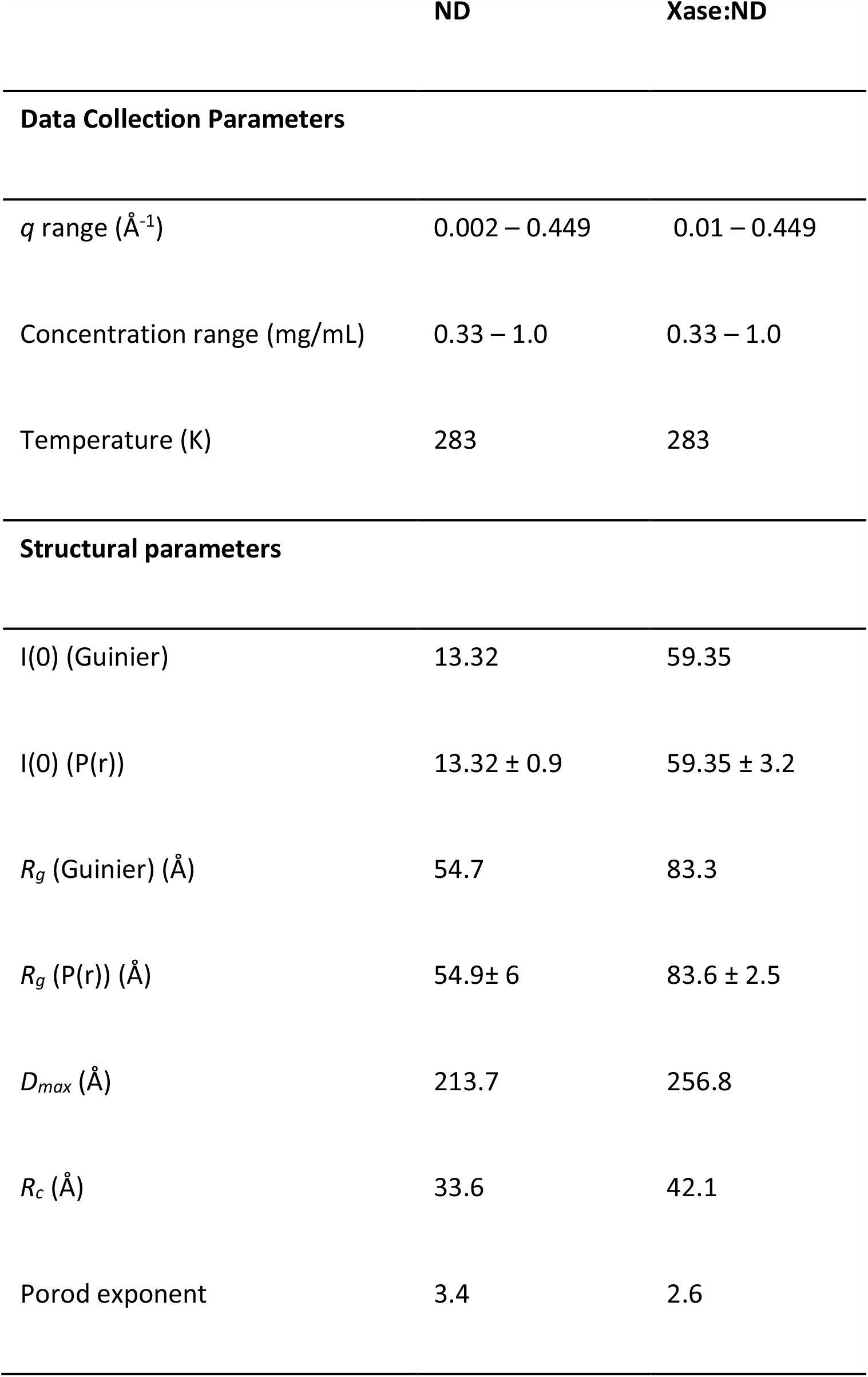
SAXS parameters for ND and Xase:ND.

|  | ND | Xase:ND |
| --- | --- | --- |
| **Data Collection Parameters** |  |  |
| $q$ range ($\text{\AA}^{-1}$) | 0.002 – 0.449 | 0.01 – 0.449 |
| Concentration range (mg/mL) | 0.33 – 1.0 | 0.33 – 1.0 |
| Temperature (K) | 283 | 283 |
| **Structural parameters** |  |  |
| $I(0)$ (Guinier) | 13.32 | 59.35 |
| $I(0)$ (P(r)) | $13.32 \pm 0.9$ | $59.35 \pm 3.2$ |
| $R_g$ (Guinier) ($\text{\AA}$) | 54.7 | 83.3 |
| $R_g$ (P(r)) ($\text{\AA}$) | $54.9 \pm 6$ | $83.6 \pm 2.5$ |
| $D_{max}$ ($\text{\AA}$) | 213.7 | 256.8 |
| $R_c$ ($\text{\AA}$) | 33.6 | 42.1 |
| Porod exponent | 3.4 | 2.6 |

**Figure 2.**
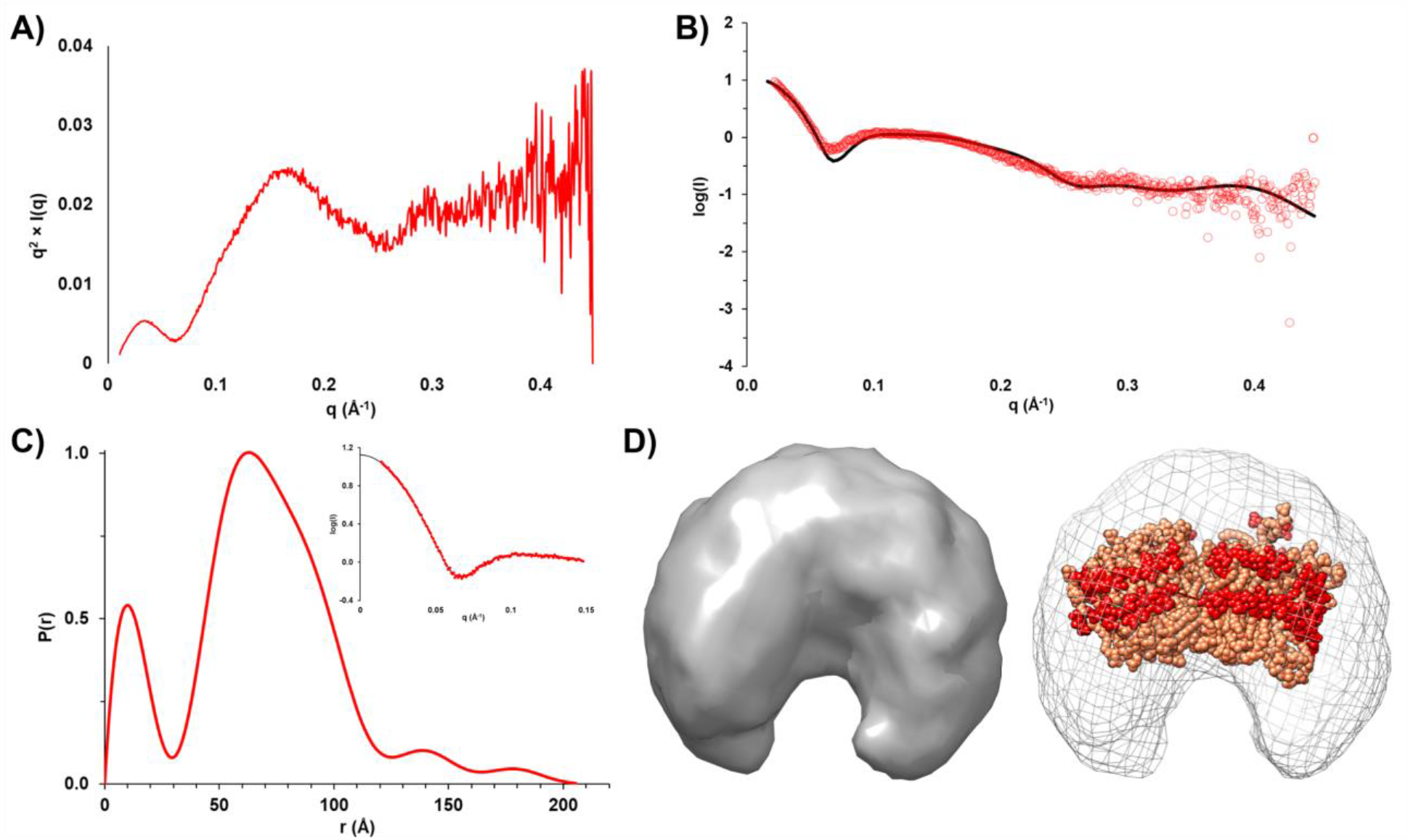
SAXS analysis of empty nanodisc. **(A)** Kratky plot analysis. **(B)** Scattering curves for experimental data (red) and theoretical model (black, χ^2^ = 0.26). **(C)** Pairwise distribution function. Inset depicts experimental data (red) and an indirect Fourier transform of the P(r) distribution plot (black), showing strong agreement between both sets of data. **(D)** (*Left*) An a*b initio* molecular envelope. (*Right*) Modeled empty nanodisc in sphere representation (scaffold protein: red, lipids: light orange) fit into the SAXS-derived molecular envelope.

### SAXS analysis of the Xase:ND complex

In order to characterize the solution structure of the Xase:ND complex, SAXS data were collected over a range of concentrations (Figure S3). Analysis of the Kratky plot confirmed that the protein sample maintained an overall globular conformation (Figure 3A). A scatter plot of the merged intensities was used to calculate a P(r) distribution plot (Figures 3B and 3C). Differences in the P(r) distribution plots between the unloaded nanodisc and Xase:ND complex indicated the two samples adopted significantly different shapes. Furthermore, calculation of the Porod exponent, a metric for the compactness of a given particle, suggested the Xase:ND adopted a more extended shape than the unloaded nanodisc (Table 2). An indirect Fourier transformation of the P(r) distribution plot was performed as a quality check, yielding a smooth fit with the experimental data in the Guinier region, supporting our calculations for a theoretical *Rg* of 83.6 ± 2.5 Å (Table 2). Because the estimated *Rc* value (42.1 Å) is approximately half of the *Rg*, we concluded that the Xase-loaded nanodisc complex adopted an elongated, cylindrical shape. A theoretical SAXS curve was calculated for the final *ab initio* envelope (Figure 3D) and aligned to our SAXS data which yielded a χ^2^ value of 0.98. The estimated *Rg* and calculated SAXS envelope (Table 2, Figure 3) were consistent with one Xase complex bound to the nanodisc, illustrating an overall elongated shape and close association between fVIIIa and fIXa (Figure 3F).

**Figure 3.**
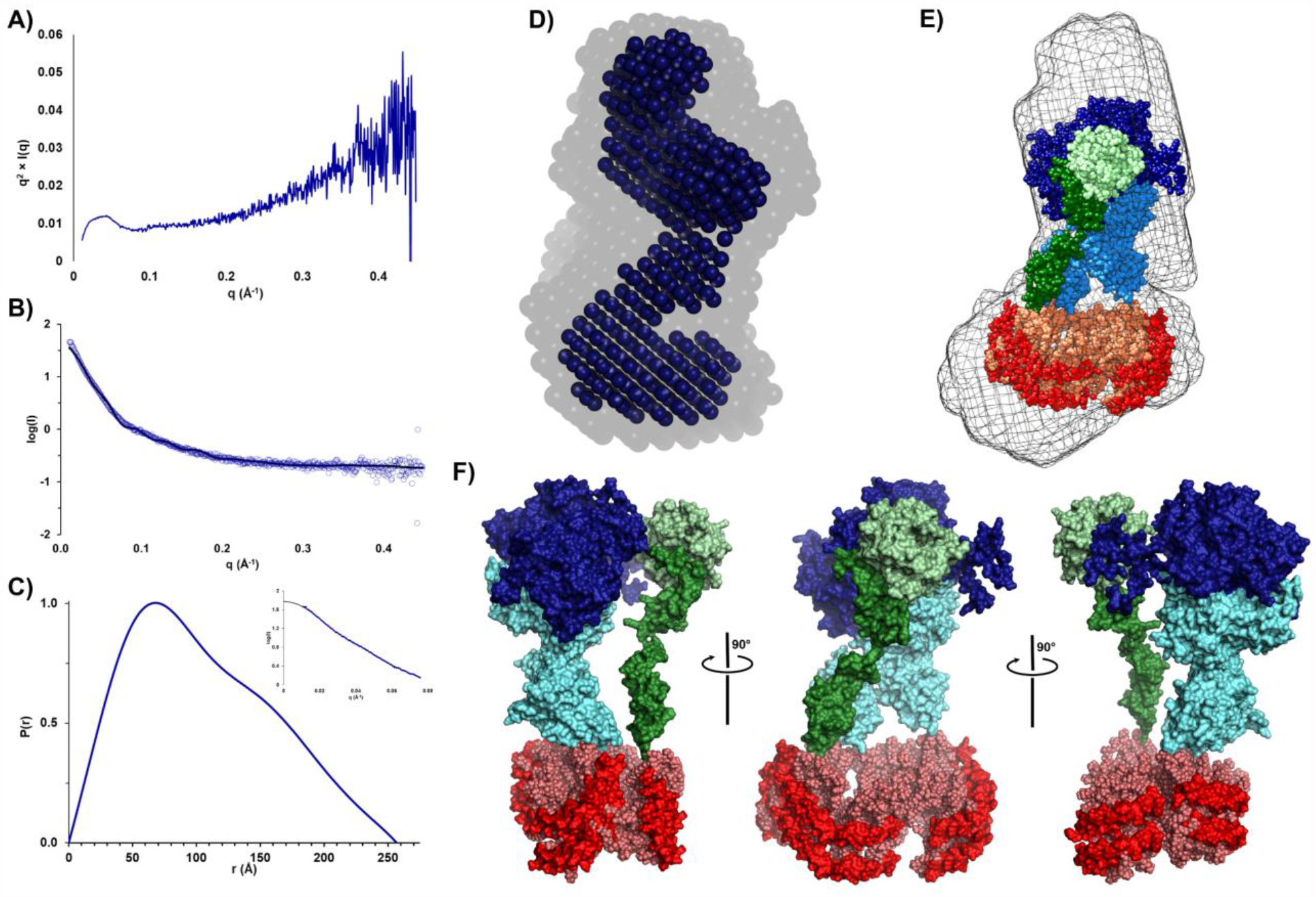
SAXS analysis of nanodisc-bound Xase complex. **(A)** Kratky plot analysis. **(B)** Scatter curves for experimental data (blue) and theoretical model (black, χ^2^ = 0.25). **(C)** Pairwise distribution function. Inset depicts experimental data (blue) and an indirect Fourier transform of the P(r) distribution plot (black), showing strong agreement between both sets of data. **(D)** *Ab initio* bead models calculated using DAMAVER (faded) and DAMFILT (solid) and aligned by SUPCOMB. **(E)** Alignment of modeled tenase complex bound to a lipid nanodisc in sphere representation (fVIIIa heavy chain: dark blue, fVIIIa light chain: cyan, fIXa heavy chain: light green, fIXa light chain: dark green, scaffold protein: red, lipids: light orange) with the calculated *ab initio* envelope (mesh) by SUPCOMB. **(F)** Model of the nanodisc-bound Xase complex (fVIIIa heavy chain: dark blue, fVIIIa light chain: cyan, fIXa heavy chain: light green, fIXa light chain: dark green, scaffold protein: red, lipids: light orange).

### Computational modeling of the Xase:ND complex

Modeling of the Xase:ND complex into the calculated SAXS envelope was accomplished through a combination of rigid body fitting and flexible refinement, guided by previously reported interactions between fVIIIa and fIXa. The A2 and A3 domains of fVIIIa, specifically the 558-loop, are predicted to be in close proximity with the fIXa catalytic domain.^28,29,33,34^ The fVIIIa C2 domain carries a putative binding site for the fIXa Gla domain,^30,31^ both of which contribute to lipid membrane binding.^7,10,18,68^ Lastly, interdomain contacts between EGF-2 and the catalytic domain of fIXa are critical for formation of the Xase complex and prevent dissociation between the fIXa heavy and light chains.^69^

Our results support multiple points of contact between fVIIIa and fIXa. The fIXa catalytic domain fits closely to the fVIIIa A2 and A3 domains. The A2 domain, specifically residues 484-489 and 555-571, is positioned near exosite II of fIXa (Figure 4A). The fIXa 99-loop, which acts as an autoinhibitory element in fIXa by blocking the nearby substrate binding pocket, is positioned near a cleft formed by fVIIIa residues 1790-1798 and 1803-1818 of the A3 domain (Figure 4B) and may be stabilized by a combination of electrostatic and hydrophobic interactions. Additionally, residues 2228-2240 of the fVIIIa C2 domain form multiple points of contact with the Gla domain of fIXa at the nanodisc surface, including residues Y1-S3, E21-F25, and V46-D47 (Figure 4C). Lastly, our analysis suggests a novel interaction between residue K80 of the fIXa EGF-1 domain and an acidic patch at the A1/A3 domain interface of fVIIIa (Figure 4D, 4E) formed by residues E144, E1969-E1970, E1827-E1828, and E1987. Comparison of our fIXa model in the Xase:ND complex with the crystal structure of porcine fIXa (PDB ID: 1PFX) indicates the Gla and EGF-1 domains undergo a ∼60° swing relative to the EGF-2 and catalytic domains, positioning residue K80 to dock with the acidic patch at the fVIIIa A1/A3 domain interface (Figure S4).

**Figure 4.**
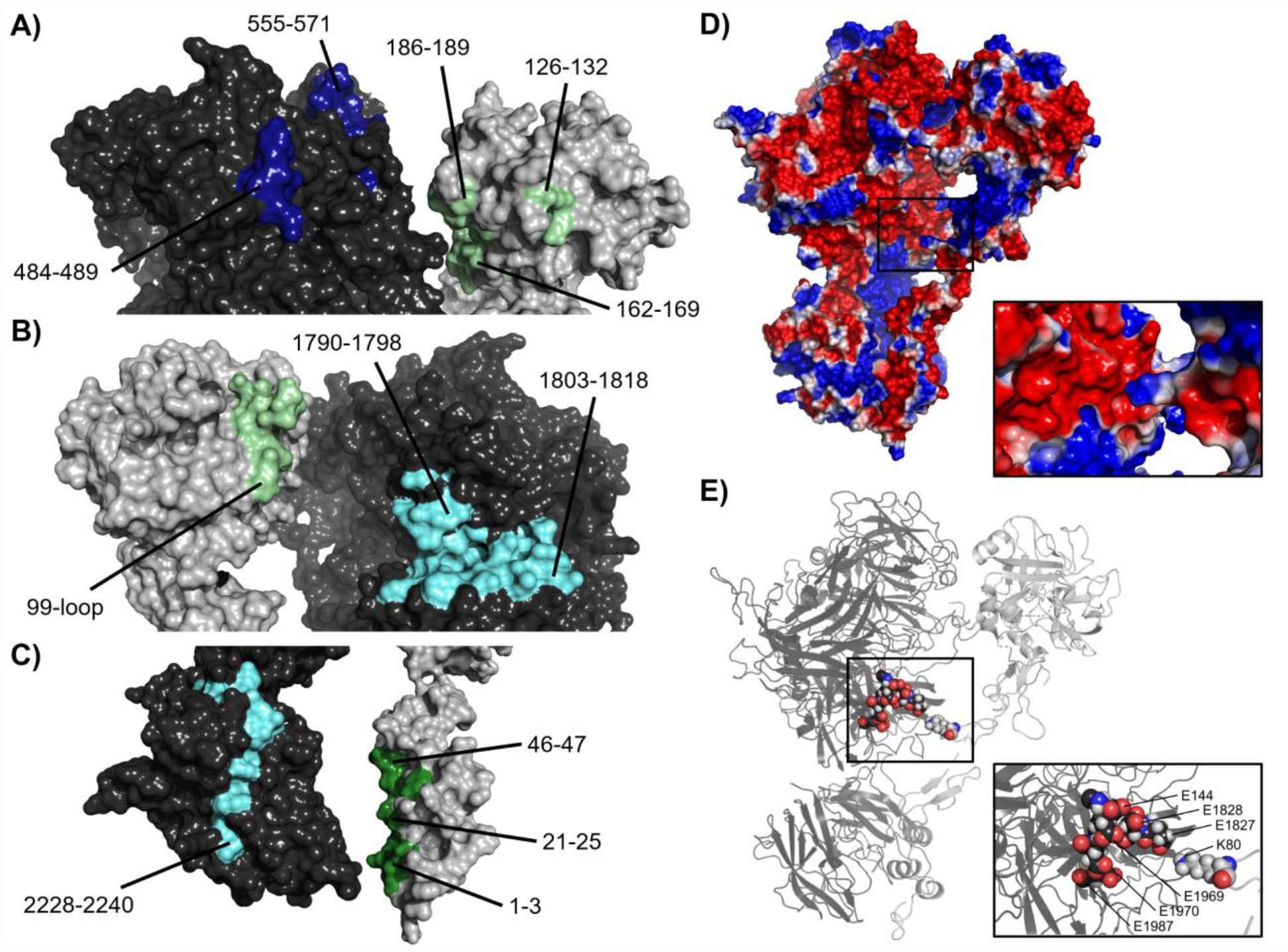
Model of Xase complex reveals multiple intermolecular contacts between fVIIIa and fIXa. **(A)** Contacts between fVIIIa A2 domain (dark blue) and fIXa exosite II (light green). **(B)** Docking between fVIIIa A3 domain (cyan) and fIXa 99-loop (light green). **(C)** Contacts between fVIIIa C2 domain (cyan) and fIXa Gla domain (dark green). **(D**,**E)** Electrostatic surface potential **(D)** and cartoon representation **(E)** of fVIIIa (dark grey) and fIXa (light grey). Inset depicts contacts between fVIIIa acidic residues and fIXa residue K80.

## Discussion

Understanding the domain organization and protein-protein interactions that mediate assembly of the lipid-bound Xase complex is of great importance for the treatment of clotting disorders. This study outlines the structural characterization of the nanodisc-bound Xase complex, revealing the domain organization and intermolecular contacts between fVIIIa and fIXa. In addition to supporting previously suggested protein-protein contacts, our analysis provides evidence for a novel interaction between the fIXa EGF-1 domain and fVIIIa A1/A3 domain interface. Our model advances our understanding of how the Xase complex is formed and how mutations at the intermolecular interface may disrupt contacts between fVIIIa and fIXa.

### Nanodiscs are a novel tool to study the intrinsic tenase complex

We calculated nanomolar affinities using His_6_-tagged nanodiscs for fVIIIa, fIXa, and the Xase complex (Table 1, Figure 1), supporting tagged nanodiscs as a convenient tool for studying lipid interactions with procoagulant proteins. Although these values are considerably higher than previously reported affinities for fVIIIa (2-5 nM), fIXa (3 nM), and Xase (0.5 nM) using activated platelets and lipid vesicles,^5,70–72^ discrepancies may be due to the reduced surface area of nanodiscs and a limited number of available binding sites. Indeed, activated platelets and lipid vesicles, prepared with 20% phosphatidylserine, present ∼600-1,200 binding sites for fVIIIa and fIXa,^72–74^ yet inactive fIX had a reported *K*_*D*_ of 390 nM with lipid nanodiscs,^75^ in good agreement with the *K*_*D*_ reported in this study for fIXa. We attempted to calculate binding affinities from a 2:1 binding model, as the nanodisc structure presents two potential binding sites on either side; however, the binding affinities were indeterminate. Furthermore, the SAXS-derived *Rg* and molecular envelope of the Xase:ND complex (Table 2, Figure 3) suggested only one Xase per nanodisc. This binding limitation may also be due to the empty nanodisc adopting a bent, elliptical conformation (Figure 2D), as previously suggested,^43,63–67^ that may promote optimal binding to one side over the other.

Previous experiments using nanodiscs^76^ and nanotubes^77^ have proposed fVIII adopts a homodimeric quaternary structure when bound to lipids. We attempted to model a fVIII homodimer using the electron crystallography-derived structure of lipid-bound fVIII (PDB ID: 3J2Q) to determine whether our data could support this model. Alignment between our experimental data and a theoretical SAXS scattering curve of dimeric fVIII bound to a nanodisc suggests poor agreement (Figure S5A), though this may be due to the presence of fIXa in our SAXS experiments which would theoretically disrupt potential fVIII dimerization.

### fVIIIa binds to fIXa via the A2, A3, and C2 domains

Several groups have provided a wealth of biochemical data on the intermolecular contacts between fVIIIa and fIXa, enhancing our understanding of how the Xase complex is formed and regulated.^27–33^ Calculating an *ab initio* molecular envelope of the Xase:ND complex allowed us to computationally dock fVIIIa to fIXa (Figure 3F) and investigate the feasibility of previously proposed protein-protein interactions as well as speculate on novel points of contact (Figure 4). Formation of the Xase complex is not predicted to alter the redox states of the disulfide bridges in fVIIIa or fIXa,^78^ limiting any conformational rearrangements to flexible linkers between domains.

First, fVIIIa residues 484-509 and 558-565 of the A2 domain and residues 1790-1798 and 1811-1818 of the A3 domain are predicted to bind to fIXa.^36,79,80^ HDX protection patterns at exosite II and the 99-loop of fIXa have recently been identified in the presence of fVIIIa and are presumed to be binding sites for the cofactor.^29,34^ Our model suggests close proximity between the fVIIIa A2-A3 domains and fIXa catalytic domain and is in good agreement with the crystal structure of the homologous prothrombinase complex.^37^ This proposed domain organization provides a rationale for how binding to fVIIIa dramatically improves fIXa activity. Our Xase model places the 99-loop near an amphipathic cleft formed by fVIIIa residues 1790-1798 and 1803-1818 of the A3 domain (Figure 4B), leading us to speculate that the fIXa 99-loop binds to fVIIIa through a combination of hydrophilic and hydrophobic interactions. Interestingly, a sequence alignment of fVIII residues 1700-1800 indicated porcine fVIII carries a glutamic acid at position 1797 while other species utilize a positively charged or neutral amino acid (Figure S6). Because the fVIII in this study is a chimera with porcine A1 and A3 domains, this extra acidic residue may strengthen interactions with the fIXa 99-loop which could partially explain why porcine fVIII has enhanced activity in the presence of fIXa.^81^ The fIXa 99-loop is also positioned near a disordered loop on the fVIIIa A2 domain, comprised of residues 721-764 which may provide additional intermolecular contacts. Although dissociation of the A2 domain during SAXS data collection was a possibility, a model of the Xase:ND complex lacking the A2 domain did not improve the fit with the experimental data (Figure S5B). Furthermore, activated ET3i, constructed with porcine A1 and A3 domains, has previously been shown to have ∼3-fold enhanced affinity for the A2 domain due to multiple stabilizing interactions at the A1/A2 and A2/A3 domain interfaces.^42,57^

Second, residues F25 and V46 of the fIXa Gla are predicted to bind to fVIIIa.^30^ Similarly, residues 2228-2240 of the fVIIIa C2 domain are a putative binding site for fIXa.^31^ Since the C1 and C2 domains of fVIIIa and the Gla domain of fIXa directly contribute to interactions with lipid membranes,^5,7,10,30,82^ we concluded that all three domains are docked onto the nanodisc surface with fVIIIa C2 domain and fIXa Gla domain in close proximity. A structure of the lipid-bound fVIII positioned the C2 domain partially embedded in the lipid membrane,^83,84^ a conformation that would prevent binding to the fIXa Gla domain.^30,31^ This C2 domain configuration remains to be supported using higher resolution techniques such as x-ray crystallography, yet may present a novel mechanism for binding to fIXa.

Third, our docking studies between fVIIIa and fIXa suggest the fIXa Gla and EGF-1 domains undergo a 60° swing relative to the fIXa crystal structure (PDB ID: 1PFX), positioning the fIXa residue K80, located in the EGF-1 domain, near the fVIIIa A1/A3 domain interface (Figures 4D and 4E). This region of fIXa has previously been shown to bind to the fVIIIa light chain in a charge-dependent manner.^85^ Mass spectrometry studies on fVIIIa indicate a potential role for this region in regulating fVIIIa stability in the presence of fIXa. Alanine substitutions to residues K1967 and K1968, located adjacent to E1969 and E1970, revealed differential roles for each amino acid residue, with K1967A diminishing fVIIIa stability and K1968A enhancing fVIIIa stability.^86^ Binding residue K80 to fVIIIa may also stabilize the fIXa EGF-1/EGF-2 interface which has been shown to modulate Xase assembly and activity.^87^ Docking the fIXa EGF-1 domain to the fVIIIa A1/A3 domain interface may be an additional protein-protein contact which stabilizes the overall Xase complex.

### Mutations at the fVIIIa/fIXa interface are associated with clotting disorders

Our model of the Xase:ND complex provided remarkable insight into how mutations at the fVIIIa/fIXa interface may inhibit formation of the complex. We analyzed the *CHAMP*/*CHBMP* hemophilia A and hemophilia B mutation databases, compiled by the Centers for Disease Control,^88^ and identified a series of mutations along the fVIIIa/fIXa interface that are associated with varying severities of hemophilia A or B (Figure 5). Intriguingly, mutations that are associated with mild-moderate hemophilia A are localized to the fVIII A1/A3 domain interface and the A3 domain near the putative binding site for the fIXa 99-loop. Severe hemophilia A mutations are localized to the A2 and C2 domains, specifically the 558-loop on the A2 domain, previously shown to be a strong determinant in binding to fIXa.^28^ We also identified several hemophilia B-related mutations in fIXa that may disrupt binding to fVIIIa, a majority of which are associated with severe cases. For instance, mutation of fIXa residue G79, located in the EGF-1 domain, to glutamic acid was identified in a hemophilia B patient with low fIXa coagulant activity.^89^ Substitution with a negatively charged amino acid would disrupt potential electrostatic contacts between fVIIIa and fIXa and reduce circulatory levels of Xase. Furthermore, residues 89-94 of the EGF-2 domain, previously shown to regulate formation and activity of the Xase complex,^69^ may play a critical role in binding to fVIIIa. Mutations to residues N92 and G93 abolished fXa generation^69,90^ and are associated with severe hemophilia B (Figure 5). Our model of the Xase complex suggests these residues form direct contacts with the fVIIIa A2 domain which may reduce the dissociation rate of the A2 domain and stabilize the active Xase complex.^91^

**Figure 5.**
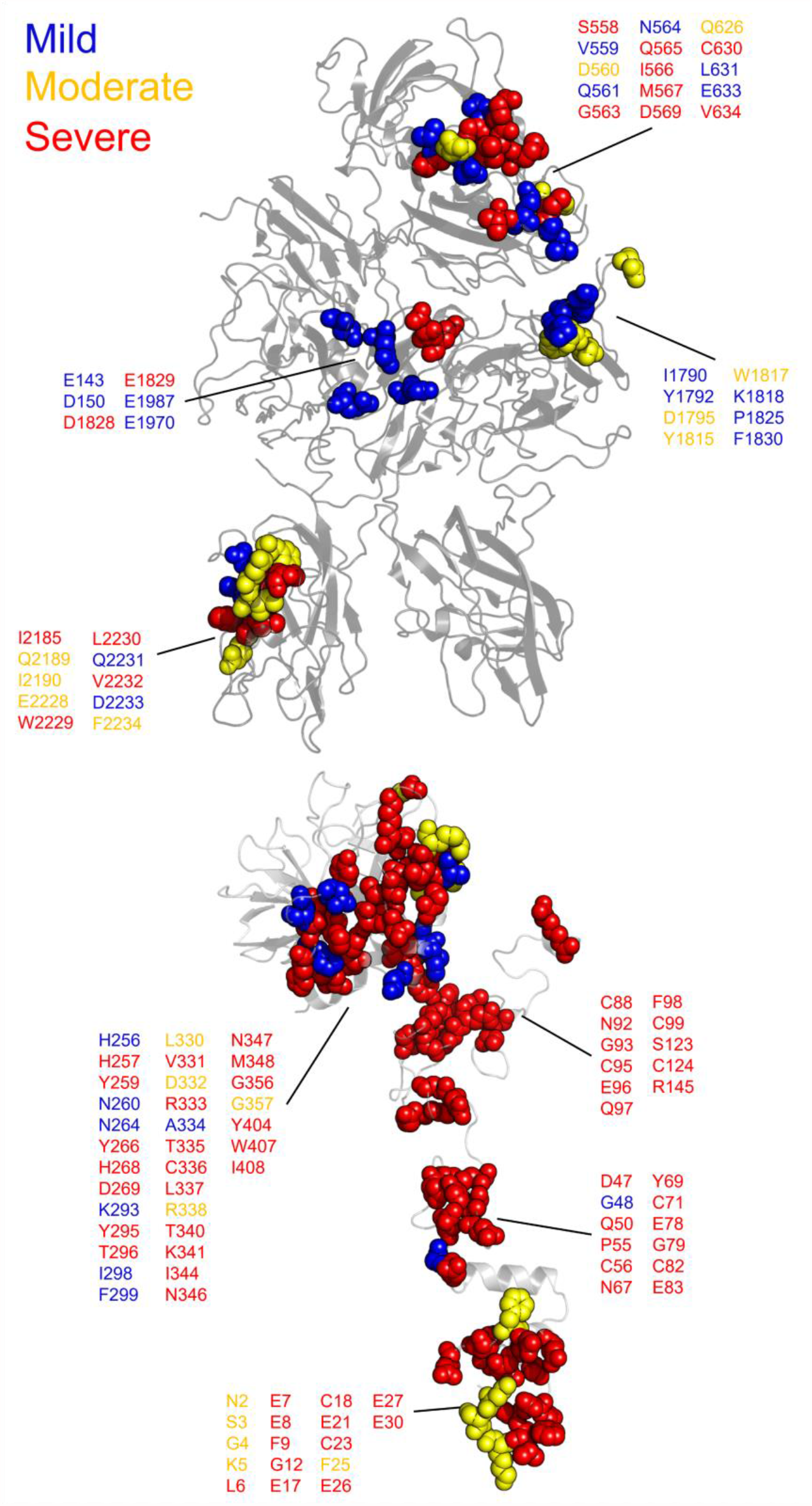
Hemophilia A and hemophilia B-related mutations to fVIIIa/fIXa dimer interface. Mutations at the fVIIIa:fIXa interface were identified from the CHAMP and CHBMP database, as collated by the Centers for Disease Control, and depicted in sphere representation on the crystal structure of human fVIII (PDB ID: 6MF2, top) for hemophilia A and SWISSMODEL of human fIXa generated from porcine fIXa (PDB ID: 1PFX, bottom, HGVS numbering) for hemophilia B. Residues are colored by disease severity (blue: mild, yellow: moderate, red: severe).

Conversely, our results also provide a mechanistic rationale for pathogenic variants on the fIXa 338-helix associated with deep vein thrombosis. Previous studies have characterized gain-of-function mutations to fIX residue R338, including the Padua mutant R338L and Shanghai mutant R338Q.^20,21^ Our modeling of the Xase complex suggests the A2 domain of fVIIIa contributes several acidic residues that may bind to and stabilize residue R338 (Figure S7), as previously proposed.^29^ Screening single-point mutations to residue R338 identified several amino acid substitutions, a majority of which were hydrophobic, that increase fIX activity, suggesting that R338 is evolutionarily conserved in order to suppress fIX activity.^22^ Hydrophobic substitutions to fIX residue R338, such as the Padua mutant, may be stabilized by fVIII residues I566 and M537. Interestingly, mutation of fVIII residues I566 or M567 to lysine or arginine is associated with moderate-severe hemophilia A,^92–94^ presumably due to charge-charge repulsion of the fIXa 338-helix. Incorporation of fIX variants V86A, located at the EGF-1/EGF-2 interface, and E277A, adjacent to the substrate binding pocket,^95^ with R338L has been shown to synergistically enhance binding to fVIIIa ∼15-fold,^96^ further demonstrating allostery between the fIX heavy and light chains in regulating Xase complex assembly and activity. A triple variant of fIX containing R318Y, R338E, and T343Y outperformed the Padua mutant as a novel gene therapeutic in treating hemophilia B with no risk of thrombophilia.^97^ Residues R338 and T343 are located at the putative fVIIIa/fIXa interface and may form stabilizing contacts with the fVIIIa A2 domain. Together, these data suggest that binding between the fVIIIa A2 domain and fIXa helix-338 is a strong determinant in formation and stabilization of the Xase complex and highlight several fIX residues to mutate in the design of next-generation hemophilia B gene therapeutics.^98^

In this study, we illustrate an updated paradigm on the membrane-bound Xase complex. Binding measurements between activated Xase proteins and immobilized nanodiscs measured nanomolar affinities. Solution scattering intensities of the Xase:ND complex allowed us to calculate biophysical parameters, such as *R*_*g*_ and *R*_*c*_, and determine an *ab initio* molecular envelope of the protease complex. Our results support multiple domain-domain contacts involved in Xase complex assembly and emphasize the detrimental effects of fVIII and fIX point mutations in the pathogenesis of blood clotting disorders.

## Acknowledgements

This work was conducted at the Advanced Light Source (ALS), a national user facility operated by Lawrence Berkeley National Laboratory on behalf of the Department of Energy, Office of Basic Energy Sciences, through the Integrated Diffraction Analysis Technologies (IDAT) program, supported by DOE Office of Biological and Environmental Research. Additional support comes from the National Institute of Health project ALS-ENABLE (P30 GM124169) and a High-End Instrumentation Grant S10OD018483. This work was supported by the Dreyfus Foundation (Henry Dreyfus Teacher-Scholar Award) and the National Institutes of Health/National Heart, Lung and Blood Institute (award numbers R15HL103518 and U54HL141981 to P.C.S., award numbers R44HL117511, R44HL110448, U54HL112309 and U54HL141981 to C.B.D., H.T.S. and P.L.).

## Author Contributions

K.C.C. planned experiments, performed experiments, analyzed data, and wrote the manuscript. S.C.P. planned experiments, performed experiments, and analyzed data. H.T.S, C.B.D. and P.L. developed expression and purification procedures for ET3i. C.B.D. and P.L. assisted in writing the manuscript. P.C.S. planned experiments, analyzed data, and wrote the manuscript.

## References

1. Fay, P. J. Factor VIII structure and function. Int. J. Hematol. 83, 103–108 (2006).

2. Bannow, B. S. et al. Factor VIII: Long-established role in haemophilia A and emerging evidence beyond haemostasis. Blood Rev. 35, 43–50 (2019).

3. Fay, P. J. Activation of factor VIII and mechanisms of cofactor action. Blood Rev. 18, 1–15 (2004).

4. Saenko, E. L. & Scandella, D. The acidic region of the factor VIII light chain and the C2 domain together form the high affinity binding site for von Willebrand factor. J. Biol. Chem. 272, 18007–18014 (1997).

5. Gilbert, G. E. & Drinkwater, D. Specific Membrane Binding of Factor VIII Is Mediated by O-Phospho-L-serine, a Moiety of Phosphatidylserine. Biochemistry 32, 9577–9585 (1993).

6. Madsen, J. J. et al. Membrane Interaction of the Factor VIIIa Discoidin Domains in Atomistic Detail. Biochemistry 54, 6123–6131 (2015).

7. Brison, C. M., Mullen, S. M., Wuerth, M. E., Podolsky, K. & Spiegel, P. C. The 1.7 Å X-Ray Crystal Structure of the Porcine Factor VIII C2 Domain and Binding Analysis to Anti-Human C2 Domain Antibodies and Phospholipid Surfaces. PLoS One 10, 1–17 (2015).

8. Brandstetter, H., Bauer, M., Huber, R., Lollar, P. & Bode, W. X-ray structure of clotting factor IXa : Active site and module structure related to Xase activity and hemophilia B. Proc. Natl. Acad. Sci. 92, 9796–9800 (1995).

9. Freedman, S. J., Furie, B. C., Furie, B. & Baleja, J. D. Structure of the Calcium Ion-Bound γ-Carboxyglutamic Acid-Rich Domain of Factor IX. Biochemistry 34, 12126–12137 (1995).

10. Muller, M. P., Wang, Y., Morrissey, J. H. & Tajkhorshid, E. Lipid specificity of the membrane binding domain of coagulation factor X. J. Thromb. Haemost. 15, 2005–2016 (2017).

11. Vadivel, K. et al. Sodium-site in serine protease domain of human coagulation factor IXa: evidence from the crystal structure and molecular dynamics simulations study. J. Thromb. Haemost. 17, 574–584 (2019).

12. Lollar, P. & Parker, C. G. pH-dependent denaturation of thrombin-activated porcine factor VIII. J. Biol. Chem. 265, 1688–1692 (1990).

13. Fay, P. J., Beattie, T. L., Regan, L. M., O’Brien, L. M. & Kaufman, R. J. Model for the factor VIIIa-dependent decay of the intrinsic factor Xase: Role of subunit dissociation and factor IXa-catalyzed proteolysis. J. Biol. Chem. 271, 6027–6032 (1996).

14. Hopfner, K. P. et al. Coagulation factor IXa: The relaxed conformation of Tyr99 blocks substrate binding. Structure 7, 989–996 (1999).

15. Zögg, T. & Brandstetter, H. Structural Basis of the Cofactor-and Substrate-Assisted Activation of Human Coagulation Factor IXa. Structure 17, 1669–1678 (2009).

16. Sichler, K. et al. Physiological fIXa Activation Involves a Cooperative Conformational Rearrangement of the 99-Loop *. J. Biol. Chem. 278, 4121–4126 (2003).

17. Rawala-Sheikh, R., Ahmad, S. S., Ashby, B. & Walsh, P. N. Kinetics of Coagulation Factor X Activation by Platelet-Bound Factor IXa. Biochemistry 29, 2606–2611 (1990).

18. Gilbert, G. E. & Arena, A. A. Activation of the factor VIIIa-factor IXa enzyme complex of blood coagulation by membranes containing phosphatidyl-L-serine. J. Biol. Chem. 271, 11120–11125 (1996).

19. Goodeve, A. C. Hemophilia B: Molecular pathogenesis and mutation analysis. J. Thromb. Haemost. 13, 1184–1195 (2015).

20. Simioni, P. et al. X-Linked Thrombophilia with a Mutant Factor IX (Factor IX Padua). N. Engl. J. Med. 361, 1671–1675 (2009).

21. Wu, W. et al. Factor IX alteration p.Arg338Gln (FIX Shanghai) potentiates FIX clotting activity and causes thrombosis. Haemotologica 106, 264–268 (2021).

22. Samelson-Jones, B. J. et al. Evolutionary insights into coagulation factor IX Padua and other high-specific-activity variants. Blood Adv. 5, 1–3 (2021).

23. Chhabra, M., Hii, Z. W. S., Rajendran, J., Ponnudurai, K. & Fan, B. E. Venous Thrombosis in Acquired Hemophilia: The Complex Management of Competing Pathologies. TH Open 03, e325–e330 (2019).

24. Weitz, J. I. Emerging anticoagulant drugs. Arterioscler. Thromb. Vasc. Biol. 27, 721 (2007).

25. Latinović, Z. et al. The First Intrinsic Tenase Complex Inhibitor with Serine Protease Structure Offers a New Perspective in Anticoagulant Therapy. Thromb. Haemost. 118, 1713–1728 (2018).

26. Zhao, L. et al. Discovery of an intrinsic tenase complex inhibitor: Pure nonasaccharide from fucosylated glycosaminoglycan. Proc. Natl. Acad. Sci. U. S. A. 112, 8284–8289 (2015).

27. Lenting, P. J., Donath, M. J. S. H., Van Mourik, J. A. & Mertens, K. Identification of a binding site for blood coagulation factor IXa on the light chain of human factor VIII. J. Biol. Chem. 269, 7150–7155 (1994).

28. Fay, P. J., Beattie, T., Huggins, C. F. & Regan, L. M. Factor VIIIa A2 subunit residues 558-565 represent a factor IXa interactive site. J. Biol. Chem. 269, 20522–20527 (1994).

29. Bajaj, S. P. et al. Factor IXa:factor VIIIa interaction: Helix 330-338 of factor IXa interacts with residues 558-565 and spatially adjacent regions of the A2 subunit of factor VIIIa. J. Biol. Chem. 276, 16302–16309 (2001).

30. Blostein, M. D., Furie, B. C., Rajotte, I. & Furie, B. The Gla domain of factor IXa binds to factor VIIIa in the tenase complex. J. Biol. Chem. 278, 31297–31302 (2003).

31. Soeda, T. et al. The Factor VIIIa C2 domain (residues 2228-2240) interacts with the factor IXa Gla domain in the factor Xase complex. J. Biol. Chem. 284, 3379–3388 (2009).

32. Griffiths, A. E., Rydkin, I. & Fay, P. J. Factor VIIIa A2 Subunit Shows a High Affinity Interaction with Factor IXa CONTRIBUTION OF A2 SUBUNIT RESIDUES 707-714 TO THE INTERACTION WITH FACTOR IXa *. J. Biol. Chem. 288, 15057–15064 (2013).

33. Fay, P. J. & Koshibu, K. The A2 subunit of factor VIIIa modulates the active site of factor IXa. J. Biol. Chem. 273, 19049–19054 (1998).

34. Freato, N. et al. Factor VIII–driven changes in activated factor IX explored by hydrogen-deuterium exchange mass spectrometry. Blood 136, 2703–2714 (2020).

35. Freato, N. et al. Probing activation-driven changes in coagulation factor IX by mass spectrometry. J. Thromb. Haemost. 1447–1459 (2021) doi:10.1111/jth.15288.

36. Lenting, P. J., Van De Loo, J. W. H. P., Donath, M. J. S. H., Van Mourik, J. A. & Mertens, K. The sequence Glu1811-Lys1818 of human blood coagulation factor VIII comprises a binding site for activated factor IX. J. Biol. Chem. 271, 1935–1940 (1996).

37. Lechtenberg, B. C. et al. Crystal structure of the prothrombinase complex from the venom of Pseudonaja textilis. Blood 122, 2777–2783 (2013).

38. Autin, L. et al. Molecular models of the procoagulant Factor VIIIa-Factor IXa complex. J. Thromb. Haemost. 3, 2044–2056 (2005).

39. Venkateswarlu, D. Structural insights into the interaction of blood coagulation co-factor VIIIa with factor IXa: A computational protein-protein docking and molecular dynamics refinement study. Biochem. Biophys. Res. Commun. 452, 408–414 (2014).

40. Ngo, J. C. K., Huang, M., Roth, D. A., Furie, B. C. & Furie, B. Crystal Structure of Human Factor VIII: Implications for the Formation of the Factor IXa-Factor VIIIa Complex. Structure 16, 597–606 (2008).

41. Doering, C. B., Healey, J. F., Parker, E. T., Barrow, R. T. & Lollar, P. Identification of Porcine Coagulation Factor VIII Domains Responsible for High Level Expression via Enhanced Secretion. J. Biol. Chem. 279, 6546–6552 (2004).

42. Parker, E. T., Doering, C. B. & Lollar, P. A1 subunit-mediated regulation of thrombin-activated factor VIII A2 subunit dissociation. J. Biol. Chem. 281, 13922–13930 (2006).

43. Denisov, I. G., Grinkova, Y. V., Lazarides, A. A. & Sligar, S. G. Directed Self-Assembly of Monodisperse Phospholipid Bilayer Nanodiscs with Controlled Size. J. Am. Chem. Soc. 126, 3477–3487 (2004).

44. Bao, H., Duong, F. & Chan, C. S. A step-by-step method for the reconstitution of an ABC transporter into nanodisc lipid particles. J. Vis. Exp. 8–13 (2012) doi:10.3791/3910.

45. Hagn, F., Nasr, M. L. & Wagner, G. Assembly of phospholipid nanodiscs of controlled size for structural studies of membrane proteins by NMR. Nat. Protoc. 13, 79–98 (2018).

46. Grushin, K., White, M. A. & Stoilova-Mcphie, S. Reversible stacking of lipid nanodiscs for structural studies of clotting factors. Nanotechnol. Rev. 6, 139–148 (2017).

47. Grinkova, Y. V., Denisov, I. G. & Sligar, S. G. Engineering extended membrane scaffold proteins for self-assembly of soluble nanoscale lipid bilayers. Protein Eng. Des. Sel. 23, 843–848 (2010).

48. Hopkins, J. B. & Thorne, R. E. Quantifying radiation damage in biomolecular small-angle X-ray scattering. J. Appl. Crystallogr. 49, 880–890 (2016).

49. Konarev, P. V., Volkov, V. V., Sokolova, A. V., Koch, M. H. J. & Svergun, D. I. PRIMUS: A Windows PC-based system for small-angle scattering data analysis. J. Appl. Crystallogr. 36, 1277–1282 (2003).

50. Svergun, D. I. Determination of the regularization parameter in indirect-transform methods using perceptual criteria. J. Appl. Crystallogr. 25, 495–503 (1992).

51. Svergun, D. I. Restoring low resolution structure of biological macromolecules from solution scattering using simulated annealing. Biophys. J. 76, 2879–2886 (1999).

52. Volkov, V. V. & Svergun, D. I. Uniqueness of ab initio shape determination in small-angle scattering Vladimir. J. Appl. Crystallogr. 36, 860–864 (2003).

53. Marcink, T. C. et al. MT1-MMP Binds Membranes by Opposite Tips of Its β Propeller to Position It for Pericellular Proteolysis. Structure 27, 281-292.e6 (2019).

54. Bibow, S. et al. Solution structure of discoidal high-density lipoprotein particles with a shortened apolipoprotein A-I. Nat. Struct. Mol. Biol. 24, 187–193 (2017).

55. Panjkovich, A. & Svergun, D. I. Deciphering conformational transitions of proteins by small angle X-ray scattering and normal mode analysis. Phys. Chem. Chem. Phys. 18, 5707–5719 (2016).

56. Waterhouse, A. et al. SWISS-MODEL: homology modelling of protein structures and complexes. Nucleic Acids Res. 46, W296–W303 (2018).

57. Smith, I. W. et al. The 3.2 Å structure of a bioengineered variant of blood coagulation factor VIII indicates two conformations of the C2 domain. J. Thromb. Haemost. 18, 57–69 (2020).

58. Lyskov, S. & Gray, J. J. The RosettaDock server for local protein-protein docking. Nucleic Acids Res. 36, 233–238 (2008).

59. Chaudhury, S. et al. Benchmarking and analysis of protein docking performance in Rosetta v3.2. PLoS One 6, (2011).

60. Lyskov, S. et al. Serverification of Molecular Modeling Applications: The Rosetta Online Server That Includes Everyone (ROSIE). PLoS One 8, 5–7 (2013).

61. Schneidman-Duhovny, D., Hammel, M., Tainer, J. A. & Sali, A. FoXS, FoXSDock and MultiFoXS: Single-state and multi-state structural modeling of proteins and their complexes based on SAXS profiles. Nucleic Acids Res. 44, W424–W429 (2016).

62. Schneidman-Duhovny, D., Hammel, M., Tainer, J. A. & Sali, A. Accurate SAXS profile computation and its assessment by contrast variation experiments. Biophys. J. 105, 962– 974 (2013).

63. Bengtsen, T. et al. Structure and dynamics of a nanodisc by integrating nmr, saxs and sans experiments with molecular dynamics simulations. Elife 9, 1–24 (2020).

64. Skar-Gislinge, N. et al. Elliptical structure of phospholipid bilayer nanodiscs encapsulated by scaffold proteins: Casting the roles of the lipids and the protein. J. Am. Chem. Soc. 132, 13713–13722 (2010).

65. Skar-Gislinge, N. et al. Small-angle scattering determination of the shape and localization of human cytochrome P450 embedded in a phospholipid nanodisc environment. Acta Crystallogr. Sect. D Biol. Crystallogr. 71, 2412–2421 (2015).

66. Denisov, I. G., McLean, M. A., Shaw, A. W., Grinkova, Y. V. & Sligar, S. G. Thermotropic phase transition in soluble nanoscale lipid bilayers. J. Phys. Chem. B 109, 15580–15588 (2005).

67. Graziano, V., Miller, L. & Yang, L. Interpretation of solution scattering data from lipid nanodiscs: J. Appl. Crystallogr. 51, 157–166 (2018).

68. Takeshima, K., Smith, C., Tait, J. & Fujikawa, K. The preparation and phospholipid binding property of the C2 domain of human factor VIII. Thromb. Haemost. 89, 788–794 (2003).

69. Fribourg, C., Meijer, A. B. & Mertens, K. The Interface between the EGF2 Domain and the Protease Domain in Blood Coagulation Factor IX Contributes to Factor VIII Binding and Factor X Activation. Biochemistry 45, 10777–10785 (2006).

70. Saenko, E. L., Scandella, D., Yakhyaev, A. V. & Greco, N. J. Activation of factor VIII by thrombin increases its affinity for binding to synthetic phospholipid membranes and activated platelets. J. Biol. Chem. 273, 27918–27926 (1998).

71. Wilkinson, F. H., Ahmad, S. S. & Walsh, P. N. The Factor IXa Second Epidermal Growth Factor (EGF2) Domain Mediates Platelet Binding and Assembly of the Factor X Activating Complex. J. Biol. Chem. 277, 5734–5741 (2002).

72. Ahmad, S. S., Rawala-Sheikh, R., Cheung, W. F., Stafford, D. W. & Walsh, P. N. The role of the first growth factor domain of human factor IXa in binding to platelets and in factor X activation. J. Biol. Chem. 267, 8571–8576 (1992).

73. Gilbert, G. E., Furie, B. C. & Furie, B. Binding of human factor VIII to phospholipid vesicles. J. Biol. Chem. 265, 815–822 (1990).

74. Ahmad, S. S., Scandura, J. M. & Walsh, P. N. Structural and functional characterization of platelet receptor-mediated factor VIII binding. J. Biol. Chem. 275, 13071–13081 (2000).

75. Medfisch, S. M., Muehl, E. M., Morrissey, J. H. & Bailey, R. C. Phosphatidylethanolamine-phosphatidylserine binding synergy of seven coagulation factors revealed using Nanodisc arrays on silicon photonic sensors. Sci. Rep. 10, 1–7 (2020).

76. Grushin, K., Miller, J., Dalm, D. & Stoilova-McPhie, S. Factor VIII organisation on nanodiscs with different lipid composition. Thromb. Haemost. 113, 741–749 (2015).

77. Dalm, D. et al. Dimeric organization of blood coagulation factor VIII bound to lipid nanotubes. Sci. Rep. 5, 1–14 (2015).

78. Cook, K. M., Butera, Diego & Hogg, P. J. Evaluation of a potential redox switch in blood coagulation tenase. Biorxiv 8, 620–628 (2019).

79. Takeyama, M., Nogami, K., Sasai, K. & Shima, M. Contribution of Factor VIII A3 Domain Residues 1793-1795 to a Factor IXa-Interactive Site. Blood 132, 1173 (2018).

80. Bloem, E., Meems, H., Mertens, K. & Meijer, A. B. A3 Domain Region 1803 – 1818 Contributes to the Stability of Activated Factor VIII and Includes a Binding Site for Activated Factor IX. J. Biol. Chem. 288, 26105–26111 (2013).

81. Doering, C. B., Healey, J. F., Parker, E. T., Barrow, R. T. & Lollar, P. High level expression of recombinant porcine coagulation factor VIII. J. Biol. Chem. 277, 38345–38349 (2002).

82. Wakabayashi, H. & Fay, P. J. Replacing thefactor VIII C1 domain with a second C2 domain reduces factor VIII stability and affinity for factor IXa. J. Biol. Chem. 288, 31289–31297 (2013).

83. Stoilova-McPhie, S., Villoutreix, B. O., Mertens, K., Kemball-Cook, G. & Holzenburg, A. 3-Dimensional structure of membrane-bound coagulation factor VIII: modeling of the factor VIII heterodimer within a 3-dimensional density map derived by electron crystallography. Blood 99, 1215–1223 (2002).

84. Stoilova-Mcphie, S., Lynch, G. C., Ludtke, S. & Pettitt, B. M. Domain organization of membrane-bound factor VIII. Biopolymers 99, 448–459 (2013).

85. Christophe, O. D., Lenting, P. J., Kolkman, J. A., Brownlee, G. G. & Mertens, K. Blood coagulation factor IX residues Glu78 and Arg94 provide a link between both epidermal growth factor-like domains that is crucial in the interaction with factor VIII light chain. J. Biol. Chem. 273, 222–227 (1998).

86. Bloem, E. et al. Mass spectrometry-assisted study reveals that lysine residues 1967 and 1968 have opposite contribution to stability of activated factor VIII. J. Biol. Chem. 287, 5775–5783 (2012).

87. Celie, P. H. N. et al. The connecting segment between both epidermal growth factor-like domains in blood coagulation factor IX contributes to stimulation by factor VIIIa and its isolated A2 domain. J. Biol. Chem. 277, 20214–20220 (2002).

88. Payne, A. B., Miller, C. H., Kelly, F. M., Michael Soucie, J. & Craig Hooper, W. The CDC Hemophilia A Mutation Project (CHAMP) Mutation List: A New Online Resource. Hum. Mutat. 34, E2382–E2392 (2013).

89. Montejo, J. M., Magallón, M., Tizzano, E. & Solera, J. Identification of twenty-one new mutations in the factor IX gene by SSCP analysis. Hum. Mutat. 13, 160–165 (1999).

90. Chang, Y.-J., Wu, H.-L., Hamaguchi, N., Hsu, Y.-C. & Lin, S.-W. Identification of Functionally Important Residues of the Epidermal Growth Factor-2 Domain of Factor IX by Alanine-scanning Mutagenesis. J. Biol. Chem. 277, 25393–25399 (2002).

91. Lamphear, B. J. & Fay, P. J. Factor IXa enhances reconstitution of factor VIIIa from isolated A2 subunit and A1/A3-C1-C2 dimer. J. Biol. Chem. 267, 3725–3730 (1992).

92. Tuddenham, E. G. D. et al. Haemophilia A: Database of ncleotide substituttions, deletions, insertions and rearrangements of the factor VIII gene. Nucleic Acids Res. 19, 4821–4833 (1991).

93. Aly, A. M. et al. Hemophilia A due to mutations that create new N-glycosylation sites. Proc. Natl. Acad. Sci. U. S. A. 89, 4933–4937 (1992).

94. Rosset, C. et al. Detection of new mutations and molecular pathology of mild and moderate haemophilia A patients from southern Brazil. Haemophilia 19, 773–781 (2013).

95. Lin, C. N. et al. Generation of a novel factor IX with augmented clotting activities in vitro and in vivo. J. Thromb. Haemost. 8, 1773–1783 (2010).

96. Kao, C. Y. et al. Incorporation of the factor IX Padua mutation into FIX-Triple improves clotting activity in vitro and in vivo. Thromb. Haemost. 110, 244–256 (2013).

97. Nair, N. et al. Gene therapy for hemophilia B using CB 2679d-GT: a novel factor IX variant with higher potency than factor IX Padua. Blood 137, 2902–2906 (2021).

98. Samelson-Jones, B. J. & Arruda, V. R. Protein-Engineered Coagulation Factors for Hemophilia Gene Therapy. Mol. Ther. - Methods Clin. Dev. 12, 184–201 (2019).

