## Supplemental Information for "SAXS analysis of intrinsic tenase complex bound to lipid nanodisc highlights intermolecular contacts between factors VIIIa/IXa"

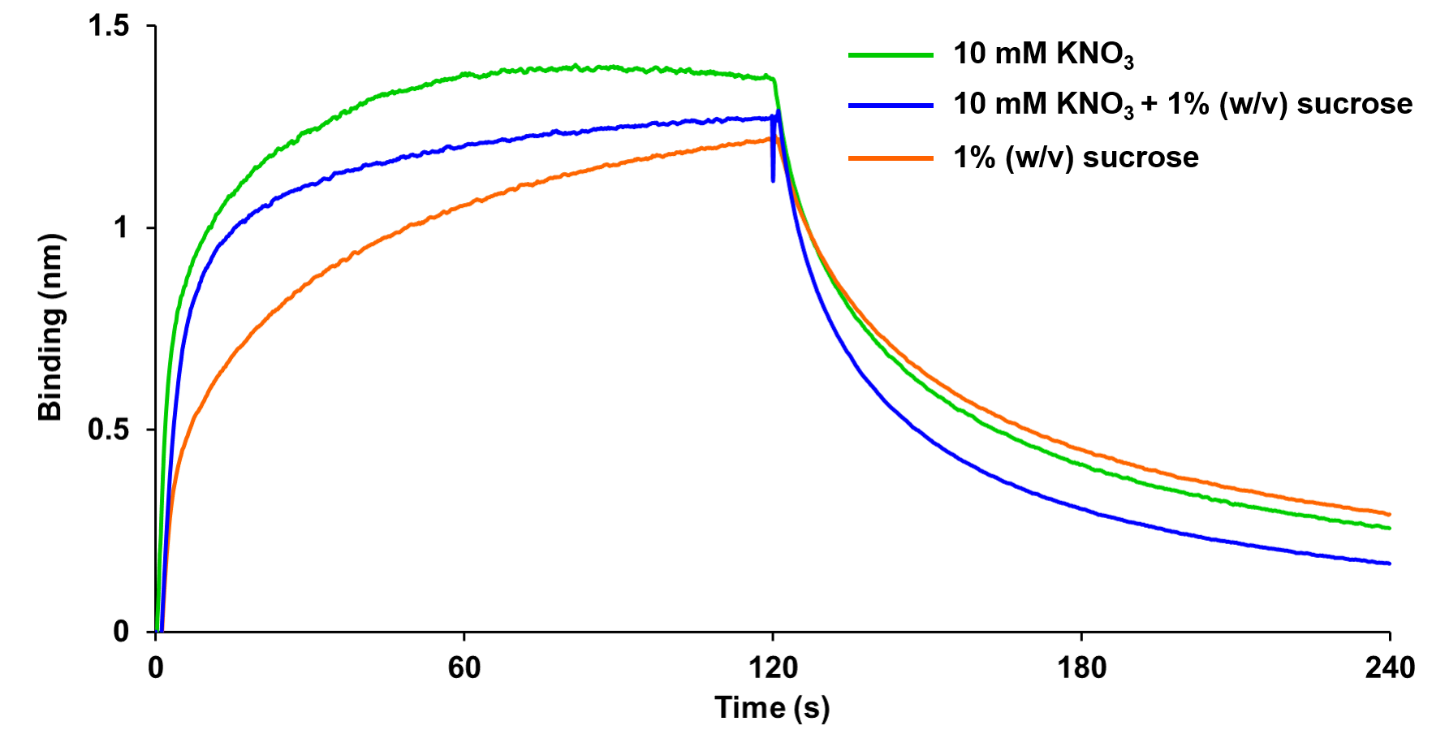

**Figure S1. Effect of radioprotective reagents on binding Xase to nanodiscs.** Binding kinetics were measured using the Xase complex (3.2 µM) and immobilized lipid nanodiscs in the presence of 10 mM KNO_3_ (green), 1% (w/v) sucrose (orange), and 10 mM KNO_3_ + 1% (w/v) sucrose (blue).

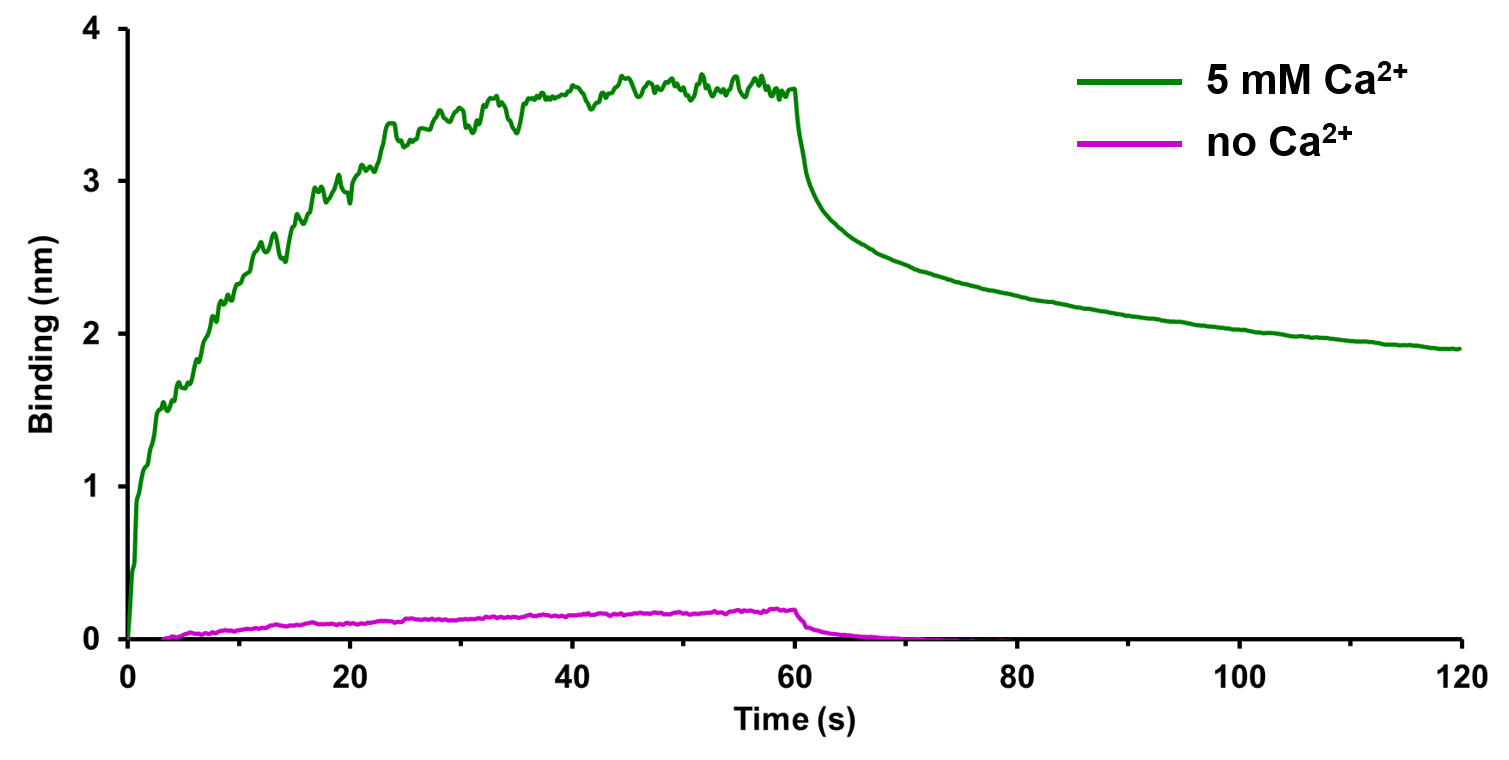

**Figure S2. Ca^2+^-dependent binding of fIXa to nanodiscs.** Bio-layer interferometry was used to determine the binding affinities of fIXa (900 nM) for immobilized nanodiscs in the absence (magenta) and presence (green) of 5 mM Ca^2+^.

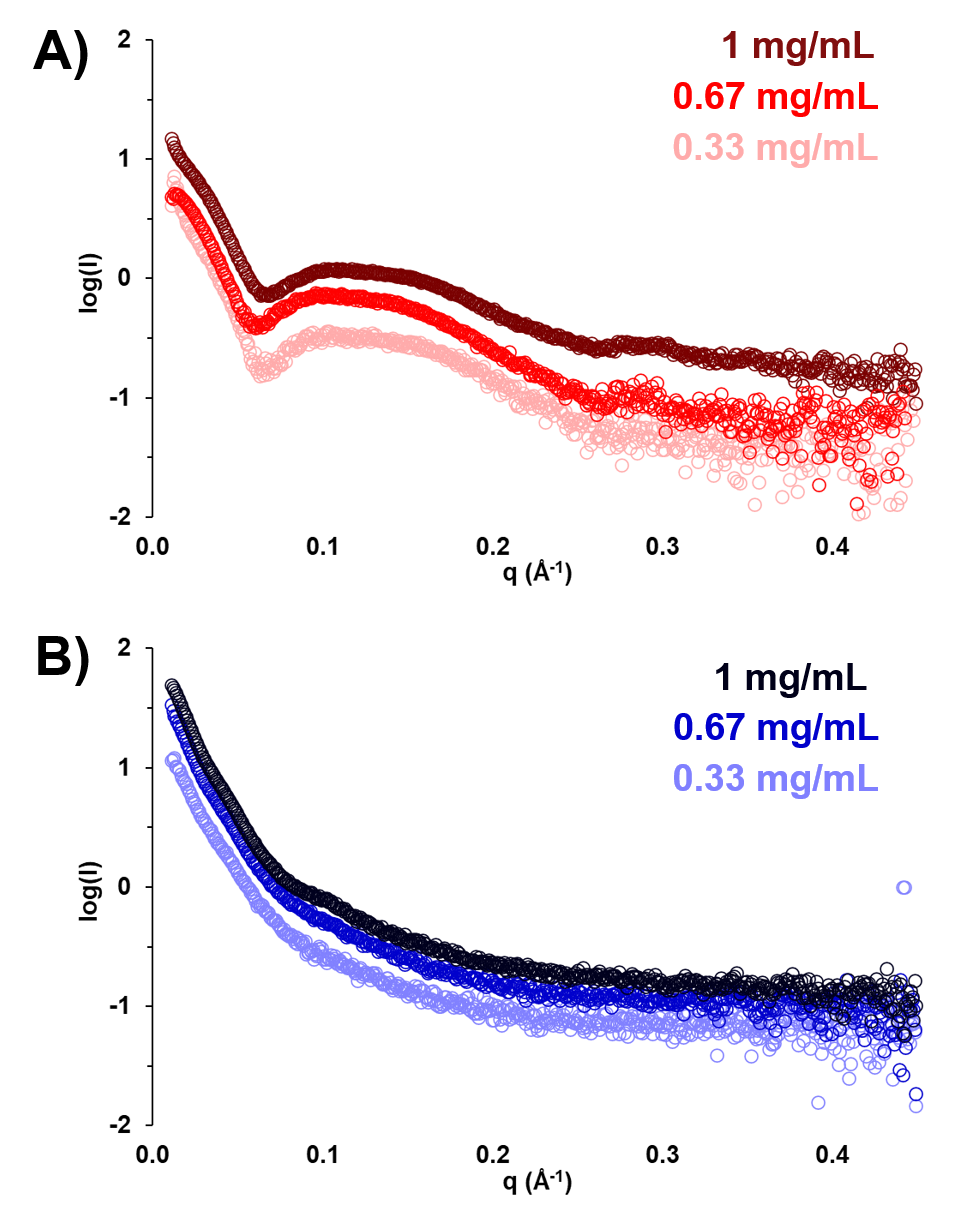

**Figure S3. Concentration-dependent SAXS scattering curves.** SAXS scattering curves were collected for the empty nanodisc (red) and nanodisc-bound Xase complex (blue) at concentrations 0.33 mg/mL (light), 0.67 mg/mL (medium), and 1 mg/mL (dark) assuming an A_280_ of 1.0 is equivalent to 1 mg/mL.

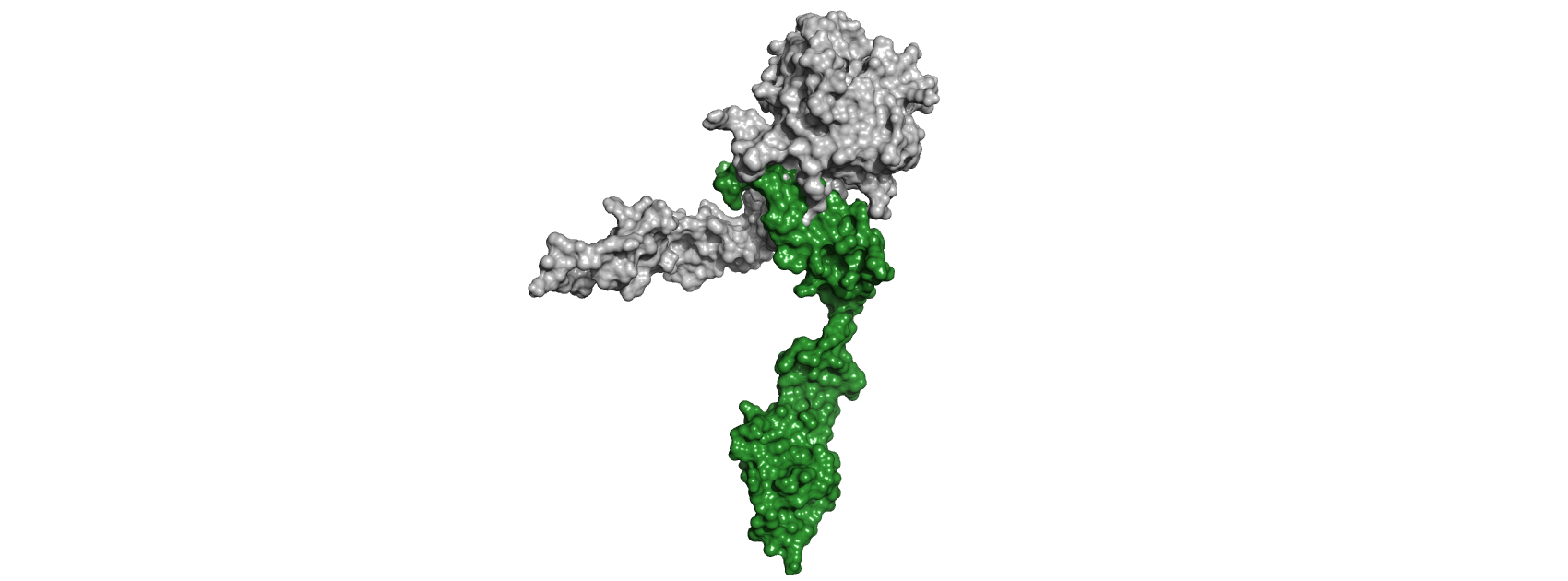

**Figure S4. Structural alignment of porcine fIXa crystal structure and modeled fIXa in Xase:ND.** Crystal structure of porcine fIXa (PDB ID: 1PFX, grey) and model of fIXa (green) in the Xase:ND complex.

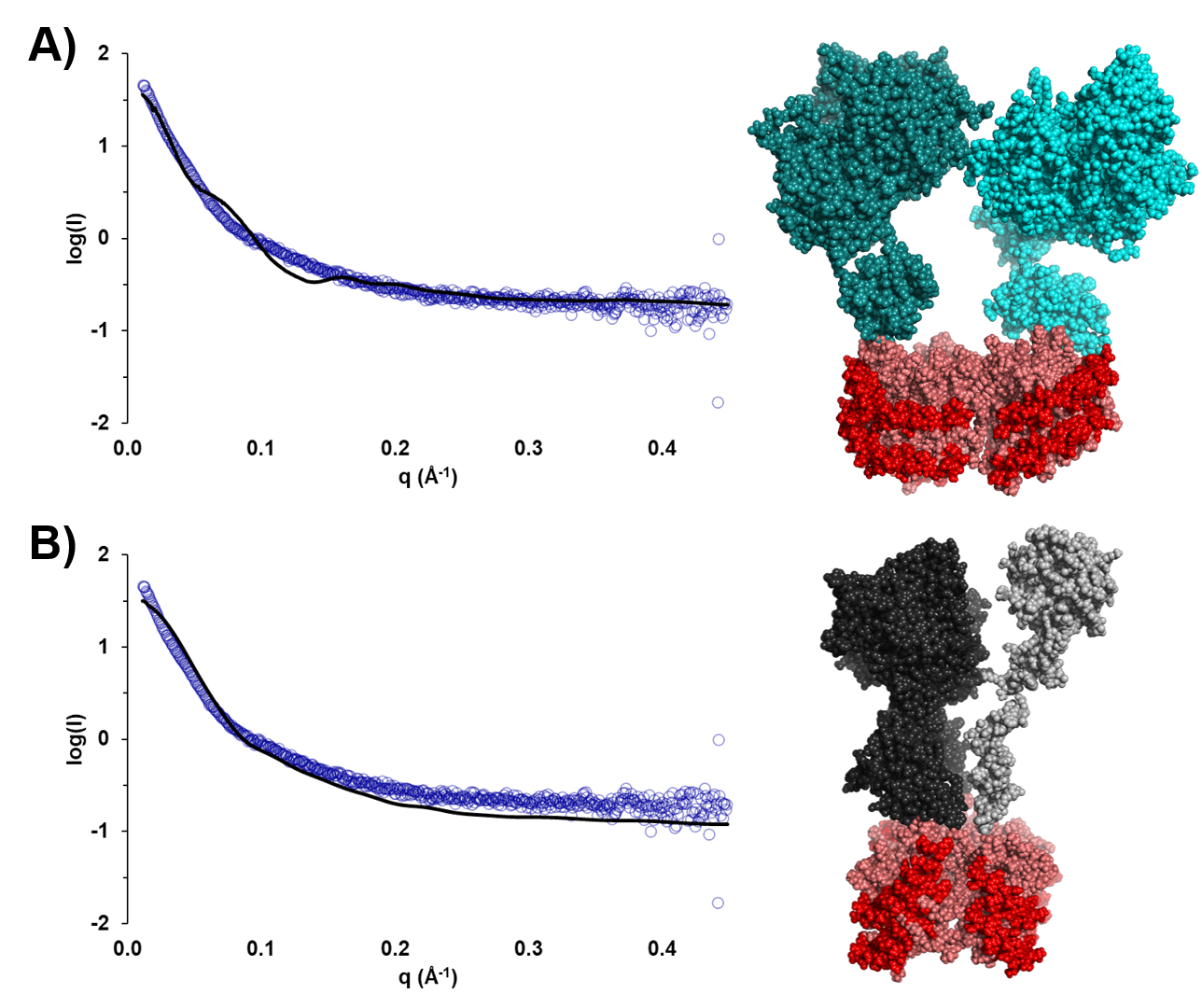

**Figure S5. Alignment of theoretical SAXS curves.** Theoretical SAXS scattering curves were calculated and aligned with experimental SAXS data (blue circles) for **(A)** homodimeric fVIII (dark/light cyan, χ^2^ = 1.40) and **(B)** Xase lacking the A2 domain (fVIIIa- dark grey, fIXa- light grey, χ^2^ = 1.04) bound to a lipid nanodisc (scaffold protein: red, lipids: light orange).

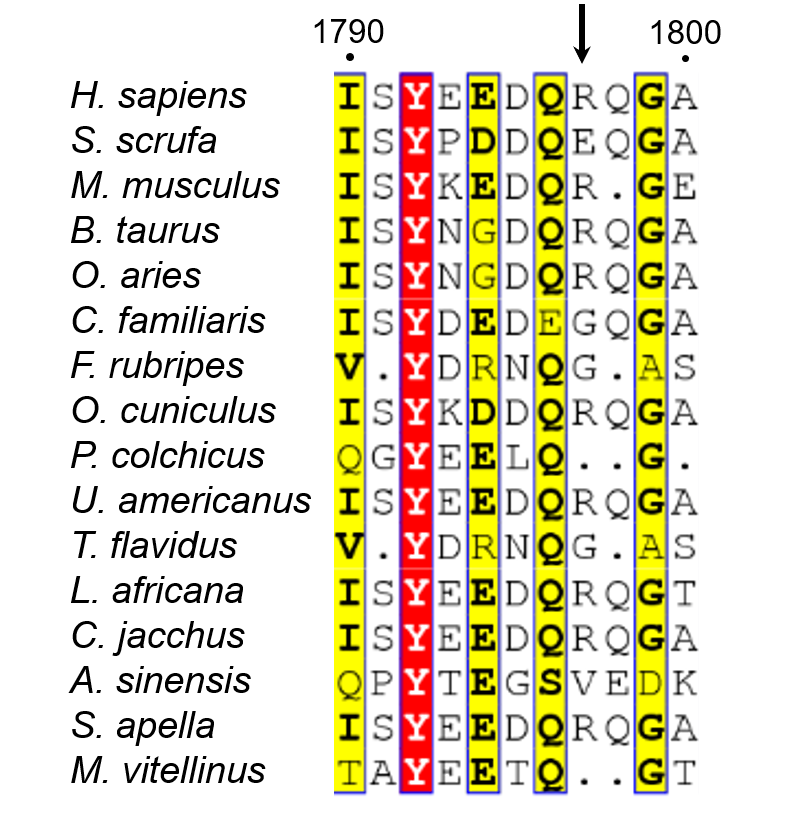

**Figure S6. Sequence alignment of fVIII residues 1790-1800.** Alignment was carried out using CLUSTAL Omega^1–3^ and visualized using ESPript.^4,5^ Identical residues are boxed in red. Similar residues are in bold and boxed in yellow. Arrow indicates variant residue 1797.

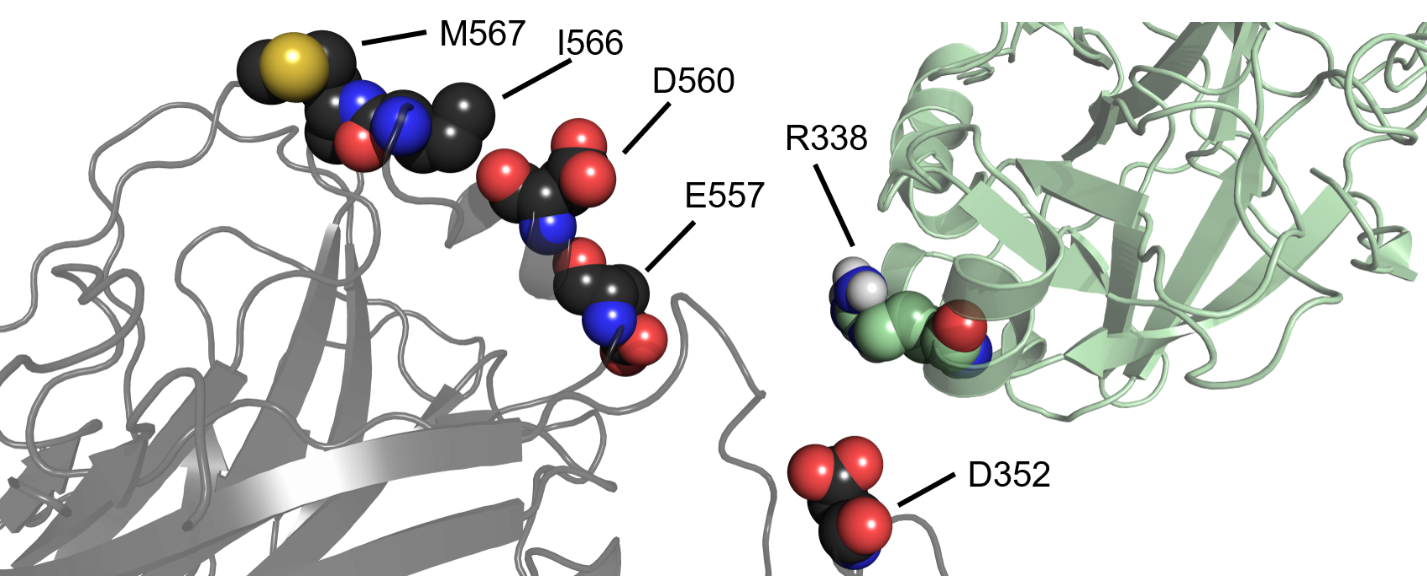

**Figure S7. Binding interface between fVIIIa A2 domain and fIXa catalytic domain.** Putative binding interface between the A2 domain of fVIIIa (dark grey) and the catalytic domain of fIXa (light green) highlights several fVIIIa residues that stabilize fIXa residue R338 (spheres).

**References**

1.	Larkin, M. A. *et al.* Clustal W and Clustal X version 2.0. *Bioinformatics* **23**, 2947–2948 (2007).

2.	Goujon, M. *et al.* A new bioinformatics analysis tools framework at EMBL–EBI. *Nucleic Acids Res.* **38**, W695-699 (2010).

3.	Chojnacki, S., Cowley, A., Lee, J., Foix, A. & Lopez, R. Programmatic access to bioinformatics tools from EMBL-EBI update: 2017. *Nucleic Acids Res.* **45**, W550–W553 (2017).

4.	Robert, X. & Gouet, P. Deciphering key features in protein structures with the new ENDscript server. *Nucleic Acids Res.* **42**, 320–324 (2014).

5.	McWilliam, H. *et al.* Analysis Tool Web Services from the EMBL-EBI. *Nucleic Acids Res.* **41**, 597–600 (2013).
